# BatchServer: a web server for batch effect evaluation, visualization and correction

**DOI:** 10.1101/2020.03.23.996264

**Authors:** Tiansheng Zhu, Guo-Bo Chen, Chunhui Yuan, Rui Sun, Fangfei Zhang, Xiao Yi, Shuigen Zhou, Tiannan Guo

**Affiliations:** Shanghai Key Lab of Intelligent Information Processing, and School of Computer Science, Fudan University, China; Zhejiang Provincial Laboratory of Life Sciences and Biomedicine, Key Laboratory of Structural Biology of Zhejiang Province, School of Life Sciences, Westlake University, 18 Shilongshan Road, Hangzhou 310024, Zhejiang Province, China; Institute of Basic Medical Sciences, Westlake Institute for Advanced Study, 18 Shilongshan Road, Hangzhou 310024, Zhejiang Province, China; Clinical Research Institute, Zhejiang Provincial People’s Hospital, People’s Hospital of Hangzhou Medical College, Hangzhou, Zhejiang, China

**Keywords:** batch effects, data pre-process, ComBat, bioinformatics, web server

## Abstract

Batch effects are unwanted data variations that may obscure biological signals, leading to bias or errors in subsequent data analyses. Effective evaluation and elimination of batch effects are necessary for omics data analysis. In order to facilitate the evaluation and correction of batch effects, here we present BatchSever, an open-source R/Shiny based user-friendly interactive graphical web platform for batch effects analysis. In BatchServer we introduced autoComBat, a modified version of ComBat, which is the most widely adopted tool for batch effect correction. BatchServer uses PVCA (Principal Variance Component Analysis) and UMAP (Manifold Approximation and Projection) for evaluation and visualizion of batch effects. We demonstate its application in multiple proteomics and transcriptomic data sets. BatchServer is provided at https://lifeinfo.shinyapps.io/batchserver/ as a web server. The source codes are freely available at https://github.com/guomics-lab/batch_server.

## INTRODUCTION

High-throughput omics data, such as mass spectrometric (MS) -based proteomics data and next generation sequencing (NGS)-based data, are usually generated with the unwanted systematic variation or so-called “batch effects”. Batch effects result from technical variations, different laboratory conditions, reagent lots, and personnels [1, 2]. If not handled properly, batch effects may distort biological signals [3] and mislead down-stream analyses [4]. Proper evaluation and correction of batch effects are thus essential in analyzing large-scale omics data.

Principal variance component analysis (PVCA) [5] and uniform manifold approximation and projection (UMAP)[6] are two widely adopted methods to identify batch effects by visualization. PVCA leverages Principal Component Analysis (PCA) and Variance Components Analysis (VCA), which fits a mixed linear model to estimate the proportion of variation of each factor. PCA [7] is a matrix factorization algorithm, for selecting top principal components as a represention of a multi-dimensional data set. VCA is a method for assessing the proportion of variance attributable to the main effects of random effect variables and the interaction of variables. PVCA combines the characteristics of both PCA and VCA, and has been applied to evaluate the batch effects and effectiveness of batch effect correction [5]. As an emerging non-linear dimensionality reduction method, UMAP is increasing used for visualization of single cell cytometry and transcriptome data [6]. We applied these two methods to evaluate batch effects.

The most intuitive way to correct batch effects is to normalize data, which adjusts the global properties of the data by comparing individual samples. However, simply exploiting normalization is sometimes insufficient to remove complex batch effects [1]. To effectively eliminate batch effects, several methods have been developed, including singular value decomposition (SVD) [8], surrogate variable analysis (SVA) [9], exploBATCH [10], BatchI [11] and ComBat [12]. Among those methods, ComBat based on a parametric or non-parametric empirical Bayes strategy is arguably the most widely method for batch correction. ComBat has been reported to be reliable and robust to outliers when tested over a large number of samples [2, 13]. However, users need to determine whether to use parametric bayes-based method or nonparametric bayes-based method. This is actually dependent on the distribution of data that is usually unknown unless evaluated [12]. Here, we modified ComBat, and implemented automated determinion of this parameter. The modified ComBat is called autoComBat. We further developed an open source web server – BatchServer that integrates PVCA, UMAP and autoComBat to provide researchers with an easy-to-use interface to evaluate and correct potential batch effects in large-scale omics data.

## MATERIALS AND METHODS

BatchServer is written in R and calls several R packages, including Shiny and Shinydashboard for R/Shiny web interface; pvca for batch effect evaluation; plotly, umap and ggplot2 for batch effect visualization; sva for batch effect correction. BatchServer is inherited from the interactive micro-service framework Shiny, integrating web interface in app.R, ui.R for the interface degisn, and server.R for data processing, other source files are called through global.R.

## RESULTS AND DISCUSSION

### The architecture design of BatchServer

To provide an easy-to-use interactive user interface for visualize the batch effects by figures and tables with option of download of the batch corrected data matrix, we designed the architecture of BatchServer as three main layers (**Figure 1**). The data input layer is provided for uploading input files of data files and sample information files and for interactively selecting batches and covariate names. The data processing layer is responsible for batch effect estimation, visualization and correction. The data output layer can display and download data processing results.

**Figure 1.**
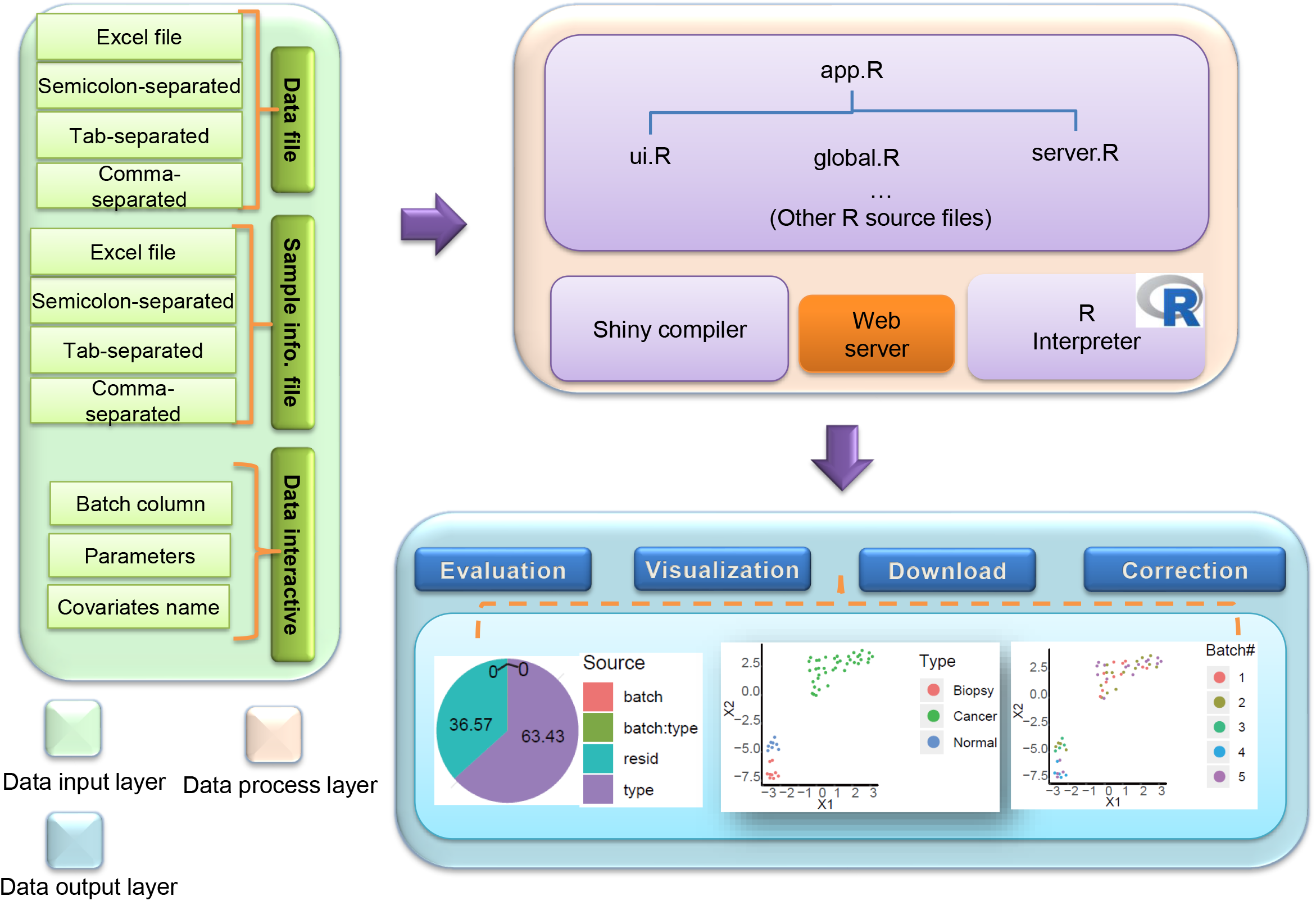
The architecture of BatchServer. Users can submit data and sample information files via the data input layer. Once submitted, files are processed by the data process layer. Thereafter, the user is required to specify potential batch effect columns and interactively set variables, if any. The data processing layer then calculates and returns the results. Finally, the data output layer will display the results interactively through figures and tables.

The detailed instructions for BatchServer are provided in the Readme section of the online web page. Here we briefly descript the usage of BatchServer. For data input, a data file and a sample information file are required. The format of these files can be tab-delimited, space-separated, comma-delimited or an Excel file. BatchServer provides a sample test data file by the bladderbatch package in the Readme section. The user can upload these two files in the “Data Input” menu, then click the “Submit” button. The data read module will read, process, and store the files for subsequent uses. Users can evaluate whether the data have batch effects using PVCA or UMAP with the online server. Both methods will display the visualization of batch effects. Once the batch effect is present, it can be adjusted using improved ComBat. Users can check and download the results of batch effect evaluation. The corrected data can be also downloaded.

### Imputation of missing values

Missing values are common in omics data sets due to technical or biological issues [14]. The mechanisms for missing values occurrence are complex and can be estimated in a number of ways [15]. Most statistical and machine learning methods do not allow a large portion of missing values; therefore, BatchServer provides four common and computationally efficient ways to replace missing values by ‘1’, ‘0’, ‘10% of the minimum’, or ‘the minimum’, where the minimum is the minimal value in the upload data matrix.

### AutoComBat: automated ComBat

Although parametric and nonparametric ComBat often brought out similar results, the parametric Bayes implantation was much computationally faster than its nonparametric one. The choice of using parametric or non-parametric Bayes implementation should be upon the goodness-of-fit test for the underlying distribution of the data set. Here we developed an automated ComBat called autoCombat in the BatchServer.

ComBat is essentially a linear model that attempts to eliminate batch effects directly from the data sets. Y_ijg_ is defined to represent the feature (e.g. protein or gene) g of sample j from batch i. Define a model that assumes

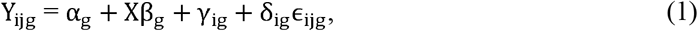

here α_g_ is the overall value of g, X is a design matrix for sample conditions, and β_g_ is the vector of the regression coefficients. The error term, ϵ_ijg_, is assumed to follow a normal distribution 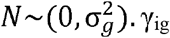 and δ_ig_ represent the additive and multiplicative batch effects of batch i for feature g, respectively. If (2) holds, ComBat will use the parametric Bayes method, otherwise it will use the non-parametric Bayes method to estimate γ_ig_ and 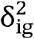.

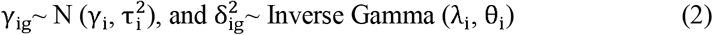

The hyperparameters γ_i_, 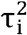, λ_i_, θ_i_ are estimated empirically from data using the method of moments[12]. When using ComBat, users must manually modify the default parameters to determine whether to use the parametric or non-parametric empirical Bayes method. For ease of use, we employed Kolmogorov-Smirnov Goodness of Fit Test (K-S) to test whether the additive parameter γ_ig_ fits normal distribution and the multiplicative parameter δ^2^_ig_ fits inverse gamma distribution for each batch at the significance level of 0.05 [16], respectively. In other words, we incorprated a ‘autoComBat” module in BatchServer which automatically switches between parametric or non-parametric Bayes method upon the detected distribution underlying the input data.

The autoComBat in BatchServer provides an additional option ‘auto’ for parameter “par.prior”, that automatically determines the use of parametric or non-parametric Bayes method. For a better visual experience, BatchServer not only plots all batches interactively, but also improves the visualization by highlighting and changing the colors of output lines and points of ComBat. Using the default parameters, we can verify the prior plot of 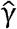 and 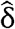 of batch ‘1’ which passed the K-S test (**Figure 2A**). Batch ‘4’ did not pass the K-S test which indicated parametric (red) and empirical (blue) estimation was not fit well (**Figure 2B**).

**Figure 2.**
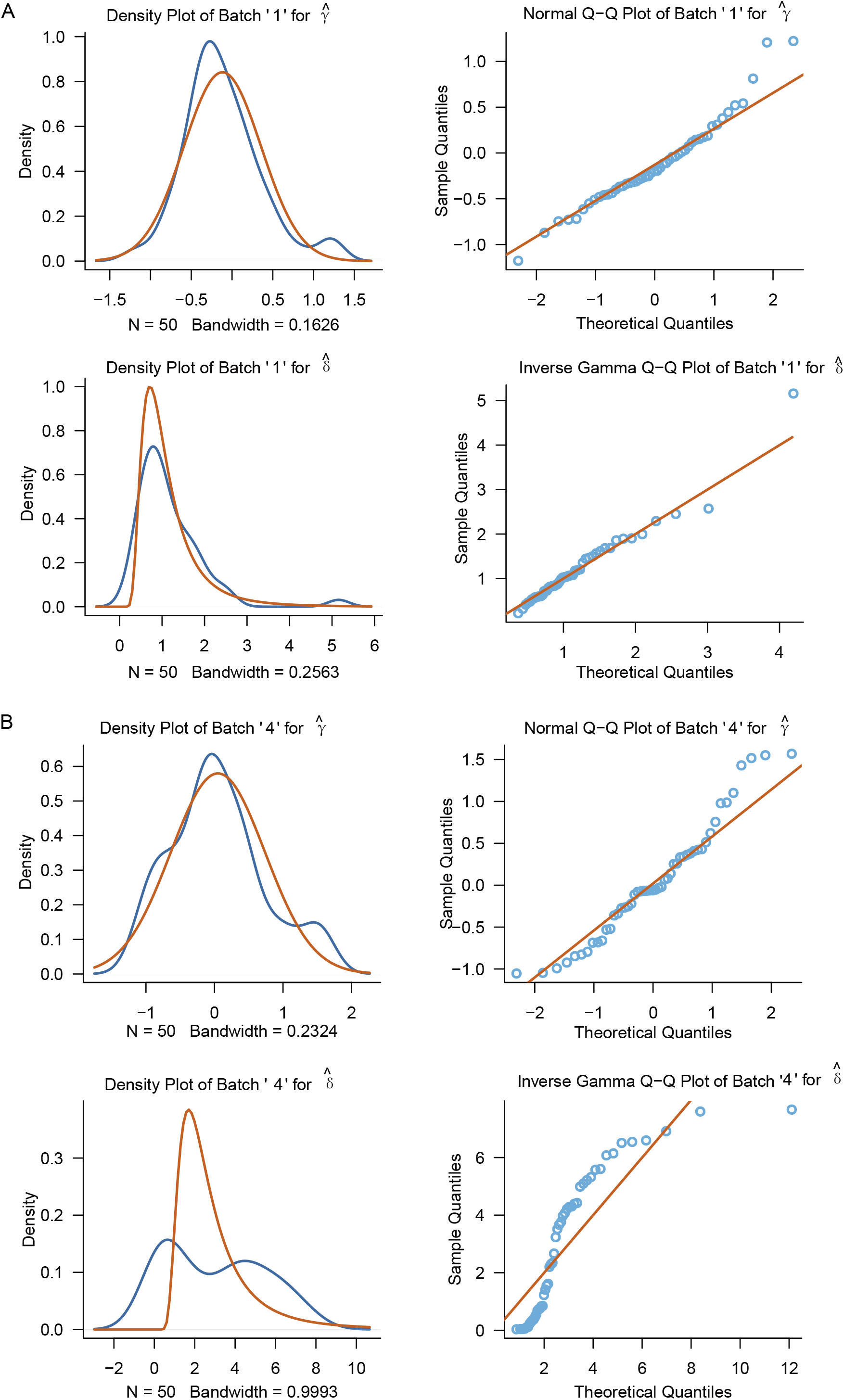
The screenshot of prior plot of γ □ and δ □ of. (A) batch ‘ 1’ which passed the K-S test and (B) batch ‘4’ which didn’t pass the K-S test using density plot and normal Q-Q plot of BatchServer. Red and blue lines/points indicate parametric and empirical estimation, respectively. The first 50 rows of bladder data in bladderbatch Bioconductor package were used as input data.

### Application of PVCA and UMAP

The PVCA method used to assess the proportion of each variable source for a given data set, but it is not universally applicable by the limitation of only accepting ExpressionSet objects. To improve the usability, we wrapped the pvcaBatchAcessess function in the pvca package and simplified the input data format to integrate the improved PVCA into BatchServer. On the other hand, UMAP was integrated into BatchServer to visualize and judge the batch effects of high dimensional omics data.

### Illustrative applications of BatchSever

To test the performance of BatchServer, we used one proteomics data set and three transcriptomics data sets for the evaluation because the latter are more accessible in the literature. As a benchmark, we used transcriptomics datasets are from the default dataset from bladderbatch package and ComBat package. We also applied the BatchServer to a published proteomic data of NCI-60 dataset cell lines [17]. The missing values of all the four data sets were replaced by ‘1’. Each data set was log2 transformed and quantile normalized as suggested by Wilson [13]. The data before and after autoComBat adjusted were analyzed and evaluated by pvca and umap module.

The first data set of transcriptomics data is a combined data matrix of GSE19804 and GSE10072 based on their shared probes, which are microarray-based transcriptomic data of 227 lung cancer disease (118 cancers and 109 controls) in female non-smokers [2]. That naturally brings out a data that were generated from two distinct batches (we named them as batch1 and batch2). **Figure 3A** showed an estimation of proportions of each factor by pvca: 1) before using autoComBat (no correction) 71.57% variation by batch, 16.87% by biological signals (tumor and normal), 0.73% to interaction between batch and biological types, and 10.84% left to residual variation; 2) after autoComBat correction, variation by batch reduced to nearly zero and consequently intensified biological variation considerably. Figure 3B and 3C showed consistent effect of batch variation correction using UMAP with Figure 3A. The results were identical when using nonparametric and auto parameters (Figure 3A, B, C) because neither batch passed the K-S test (Figure 3D). Although the computational cost for method choice was negligible for ComBat (Supplementary Table 1), the nonparametric estimation was always computational expensive than the parametric one. We also tested another two transcriptomics datasets. Both showed substantial batch effects as visualized by our BatchServer, but after applying automated procedures the batch effects were well controlled, leading to enhanced biological signals. Details were presented in Supporting Information.

**Figure 3.**
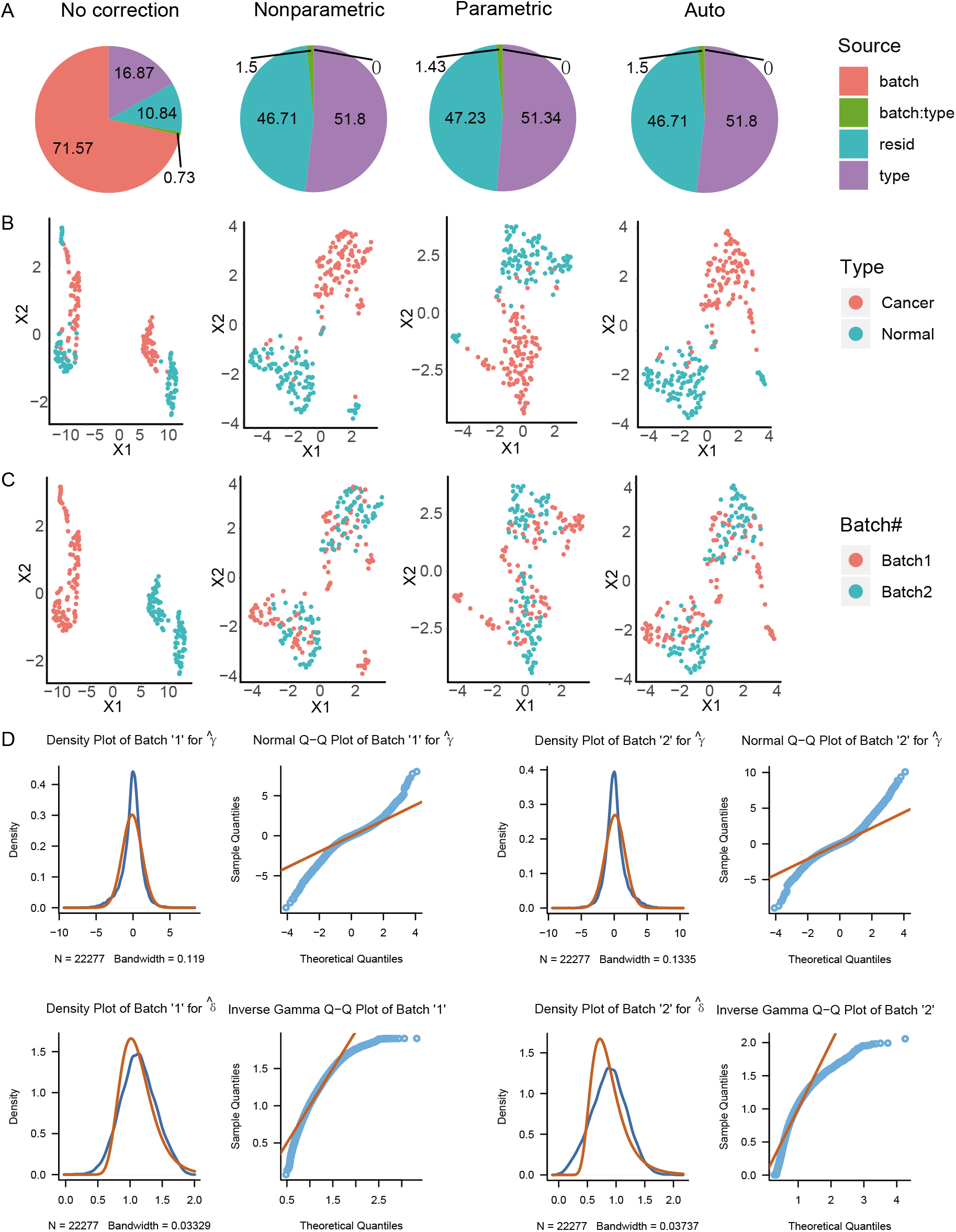
Improved performance of ComBat by the first transcriptomics data set. (A) Pie plots of batch effect using PVCA with no correction and par.prior set to nonparametric, parametric or auto for autoComBat. The number in each pie chart is the percentage of the source of variation corresponding to the respective color. ‘type’ means the variation caused by the tumor type. ‘batch’ means the variation caused by the batch number. ‘batch:type’ means interaction effects and ‘resid’ represent residual variance (unexplained variance by the chosen effects in the model, which may be caused by individual heterogeneous and other factors). (B) UMAP plots show the clustering of (B) biological interesting type and (C) batch# before and after batch effect correction by parametric ComBat, non-parametric ComBat and autoComBat, respectively. (D) Priori plots of batch effect by autoComBat.

The proteomics data set are from NCI-60 cell lines[17], containing 3,171 proteins and 60 samples of 12 batches. This data set was generated by Pressure Cyling Technology coupled with data independent acquisition (PCT-DIA)[18]. The potential factor of batch effects could be from the different tumor type or from PCT. We processed this data set with the same procedure to show the performance of batch effect evaluation and correction of BatchServer. The results are shown in **Figure 4**. The batch accounted for 3.02% variability in the data, while biological type captures 33.74% variability (Figure 4A column 1). After batch effect correction, biological type variability were enhanced (larger than 45%) and percent of batch effect fell to 0 (figure 4A column 2 to 4). That gives consistent results using UMAP (Figure 4BC), which shows batch effects were further reduced and ability to distinguish between tumor types were increased. This suggests that the batch effect of proteome data can be effectively adjusted by BatchServer.

**Figure 4.**
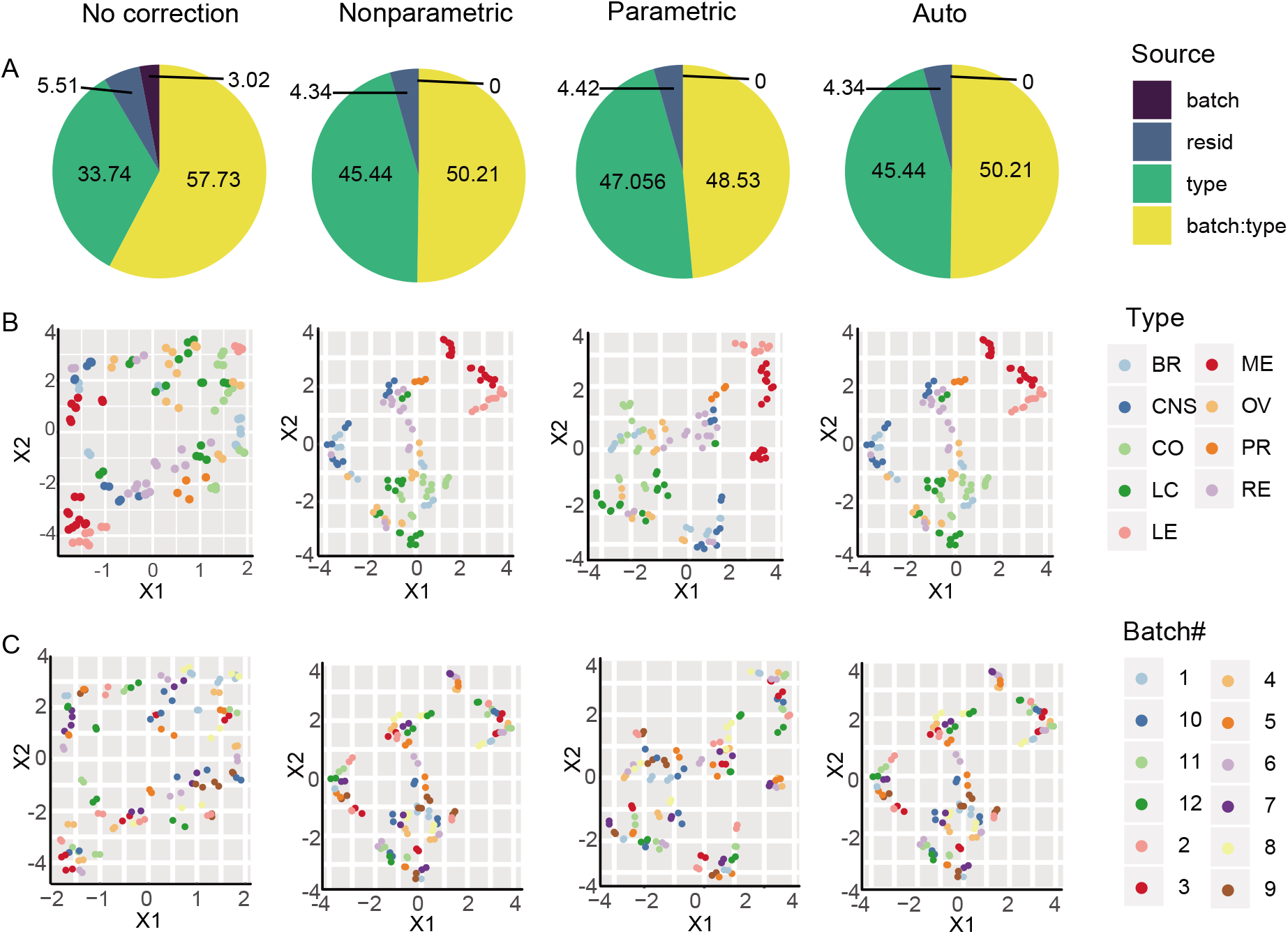
Performance of BatchServer by proteomics data set. The four columns are, in order, original data with no correction, corrected data with ‘nonparametric’, corrected data with ‘parametric’ and correct data with ‘automatic’ by autoCombat. (A) Pie plots of batch effect using PVCA. (B) UMAP visualization of the clustering of biological effect enhanced after batch effect correction. (C) UMAP show the degree of accumulation and dispersion of batches before and after batch effect correction.

## CONCLUSIONS

We developed a web server called BatchServer to facilitate the evaluation, visualization, and correction of batch processing results for large-scale omics data sets. The automated ComBat can automatically select parametric or non-parametric empirical Bayes methods for batch calibration. It also integrates PVCA and UMAP to evaluate and visualize potential batch effects. BatchServer has an R/Shiny graphical user interface for enhanced usability and is easy to install on a personal computer or server, thus providing a convenient service to process batch effect for the community. It can be effectively applied to multiple omics data sets.

## ASSOCIATED CONTENT

### Supporting Information

The following files are available free of charge.

The second and the third transcriptomic data sets description

Supplementary Figure 1-3. (file type docx)

Supplementary Figure 1: Performance of improved ComBat by the second transcriptomic data set.

Supplementary Figure 2: Visualization of BatchServer by the third transcriptomic data set.

Supplementary Figure 3: Prior plot of batch fitting effect by improved ComBat by the third transcriptomic data set.

Supplementary Table 1-3. (file type docx)

Supplementary Table 1: Time consumed (seconds) for batch effect adjust using ComBat in transcriptomic data set 1.

Supplementary Table 2: Time consumed (seconds) for batch effect adjust using improved ComBat in transcriptomic data set two.

Supplementary Table 3: Time consumed (seconds) for batch effect adjust using ComBat in transcriptomic data set 3.

## AVAILABILITY OF DATA AND MATERIALS

The proposed BatchServer and its manual are deployed at https://lifeinfo.shinyapps.io/batchserver/ and tested with Chrome, Firefox and IE browsers. Source codes and testing data are freely available at https://github.com/guomics-lab/batch_server with license for freely available to academic researchers.

## ABBREVIATIONS

MS: mass spectrometry
NGS: next generation sequencing
PCA: principal components analysis
PVCA: Principal Variance Component Analysis
UMAP: Manifold Approximation and Projection
SVD: Singular Value Decomposition
t-SNE: t-distributed stochastic neighbor embedding
K-S test: Kolmogorov–Smirnov test
DIA: data independent acquisition.

## ACKNOWLEDGMENTS

This work was supported by National Natural Science Foundation of China (General Program) (Grant No. 81972492 to T.G.), Zhejiang Provincial Natural Science Foundation for Distinguished Young Scholars (Grant No. LR19C050001 to T.G.), Hangzhou Agriculture and Society Advancement Program (Grant No. 20190101A04 to T.G.).

## AUTHORS’ CONTRIBUTIONS

TZ design and implemented the web application; TG, SZ, and TZ wrote the manuscript and conceived the project; GC, CY, RS and FZ revised and modified the manuscript and provided many useful suggestions. RS and XY conducted the software testing. All authors have read and approved the final manuscript.

## For TOC only

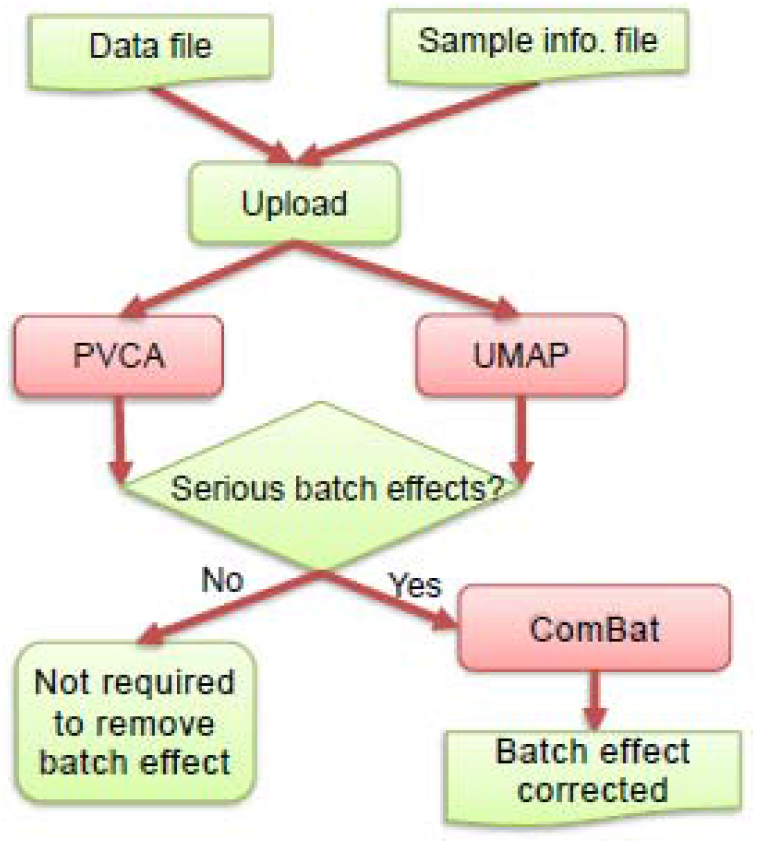

