## Supporting Information for "BatchServer: a web server for batch effect evaluation, visualization and correction"

**The second and the third transcriptomic data sets**

The second transcriptomic data set came from Bioconductor package BatchQC sample data, which captured 89 samples when activating 9 different growth pathway genes (1,600 genes) in human mammary epithelial cells (GEO accession: GSE73628). The data contained three batches and ten different conditions. We did the same experiment on this data set as transcriptomic data set one. The performance of BatchServer were similar to first data set, and details in Supplementary Table 2 and Supplementary Figure 1.

The third transcriptomic data set was from Bioconductor library bladderbatch, a microarray gene expression data with 52 genes on 57 bladder samples in 5 batches. The time cost was presented in Supplementary Table 3. The results were also very similar to the first transcriptomic data set Supplementary Figure 2ABC. It was obviously to see from Supplementary Figure 3 that when $\hat{\delta}$ was poorly fit non-parametric Bayes implementation of ComBat was in use.

The experiment of the three data sets indicate BatchServer were suitable for evaluating and removing batch effects and enhance biological signals.

**Supplementary Table 1.**

| Parameter | User | System | Elapsed |
| --- | --- | --- | --- |
| nonparametric | 3195.69 | 203.70 | 3400.65 |
| parametric | 0.53 | 0.11 | 0.64 |
| auto | 3209.11 | 239.84 | 3452.00 |

**Supplementary Table 1. Time consumed (seconds) for batch effect adjust using ComBat in transcriptomic data set 1.** The ‘User’ time is the CPU time charged for the execution of user instructions of the calling process. The ‘System’ time is the CPU time charged for execution by the system on behalf of the calling process, and the ‘Elapsed’ time is the ‘real’ elapsed time since the process was started (similarly hereinafter).

**Supplementary Table 2.**

| Parameter | User | System | Elapsed |
| --- | --- | --- | --- |
| nonparametric | 19.16 | 0.10 | 19.25 |
| parametric | 0.05 | 0.00 | 0.04 |
| auto | 18.20 | 0.25 | 18.50 |

**Supplementary Table 2. Time consumed (seconds) for batch effect adjust using autoComBat in transcriptomic data set two.**

**Supplementary Table 3**

| parameter | user | system | elapsed |
| --- | --- | --- | --- |
| parametric | 0.26 | 0.00 | 0.26 |
| nonparametric | 6483.32 | 0.46 | 6488.14 |
| auto | 6577.83 | 0.76 | 6580.92 |

**Supplementary Table 3. Time consumed (seconds) for batch effect adjust using ComBat in transcriptomic data set 3**.

**Supplementary Figure 1**
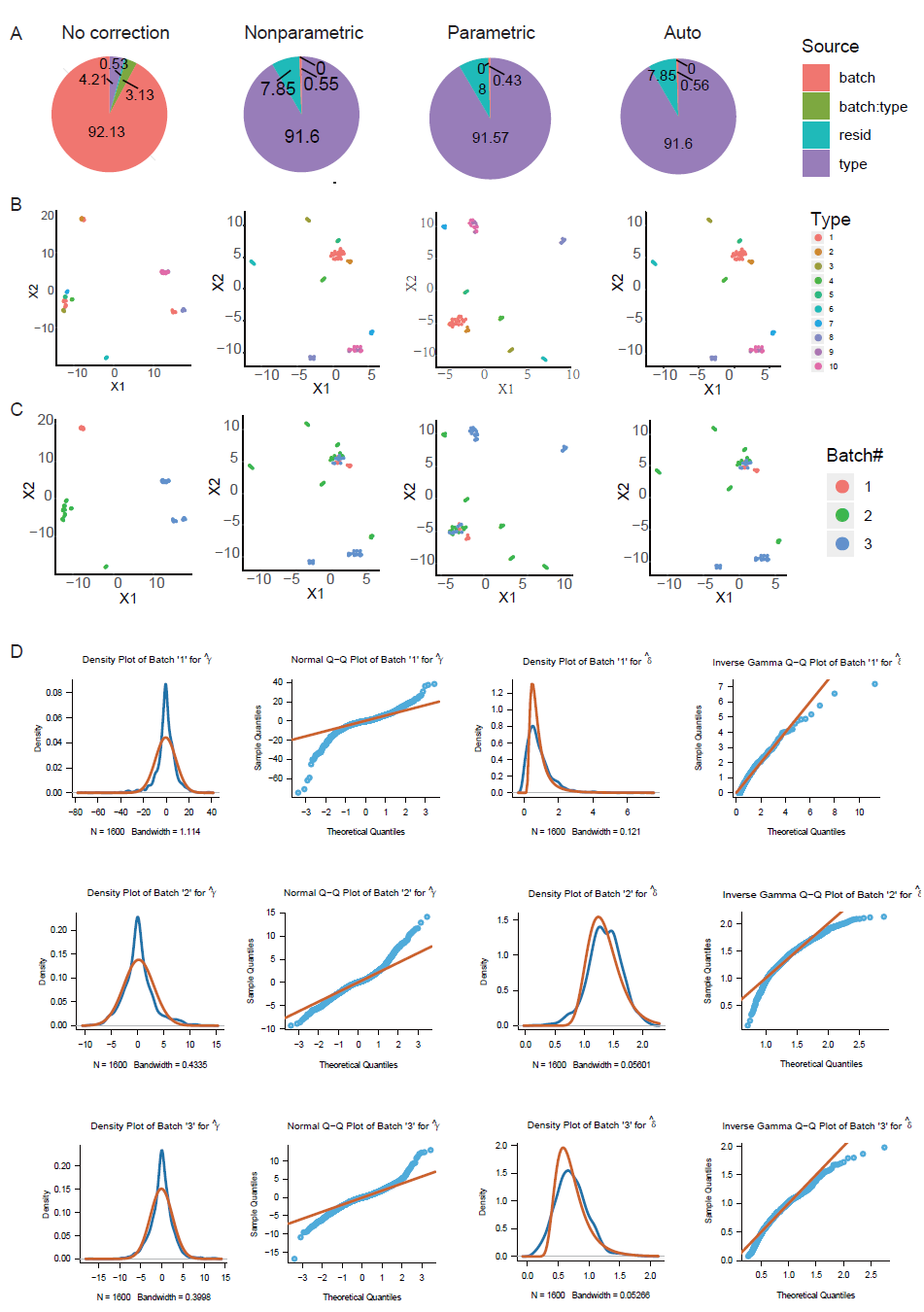

**Supplementary Figure 1. Performance of autoComBat by the second transcriptomic data set.** A) Pie plots of batch effect using PVCA with no correction and par.prior set to nonparametric, parametric or auto for autoComBat. BC) UMAP plots show the biological and batch effect clustering, respectively. D) Prior plots of batch fitting effect by autoComBat.

**Supplementary Figure 2**

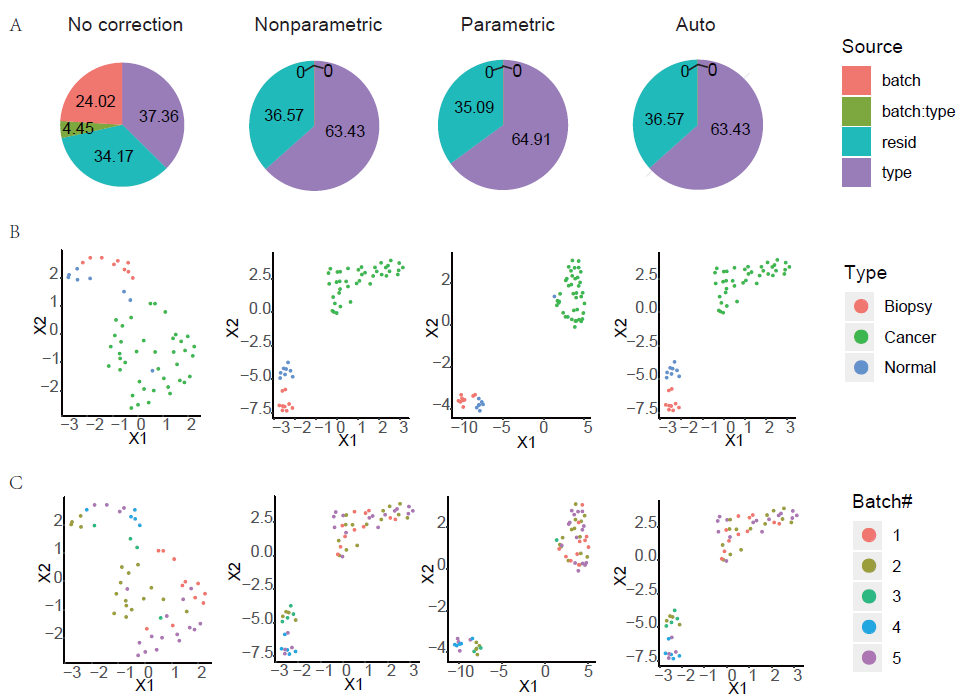

**Supplementary Figure 2. Performance of autoComBat by the third transcriptomic data set.** A) Pie plots of batch effect using PVCA with no correction and par.prior set to nonparametric, parametric or auto for autoComBat. BC) UMAP plots show the biological and batch effect clustering, respectively.

**Supplementary Figure 3**

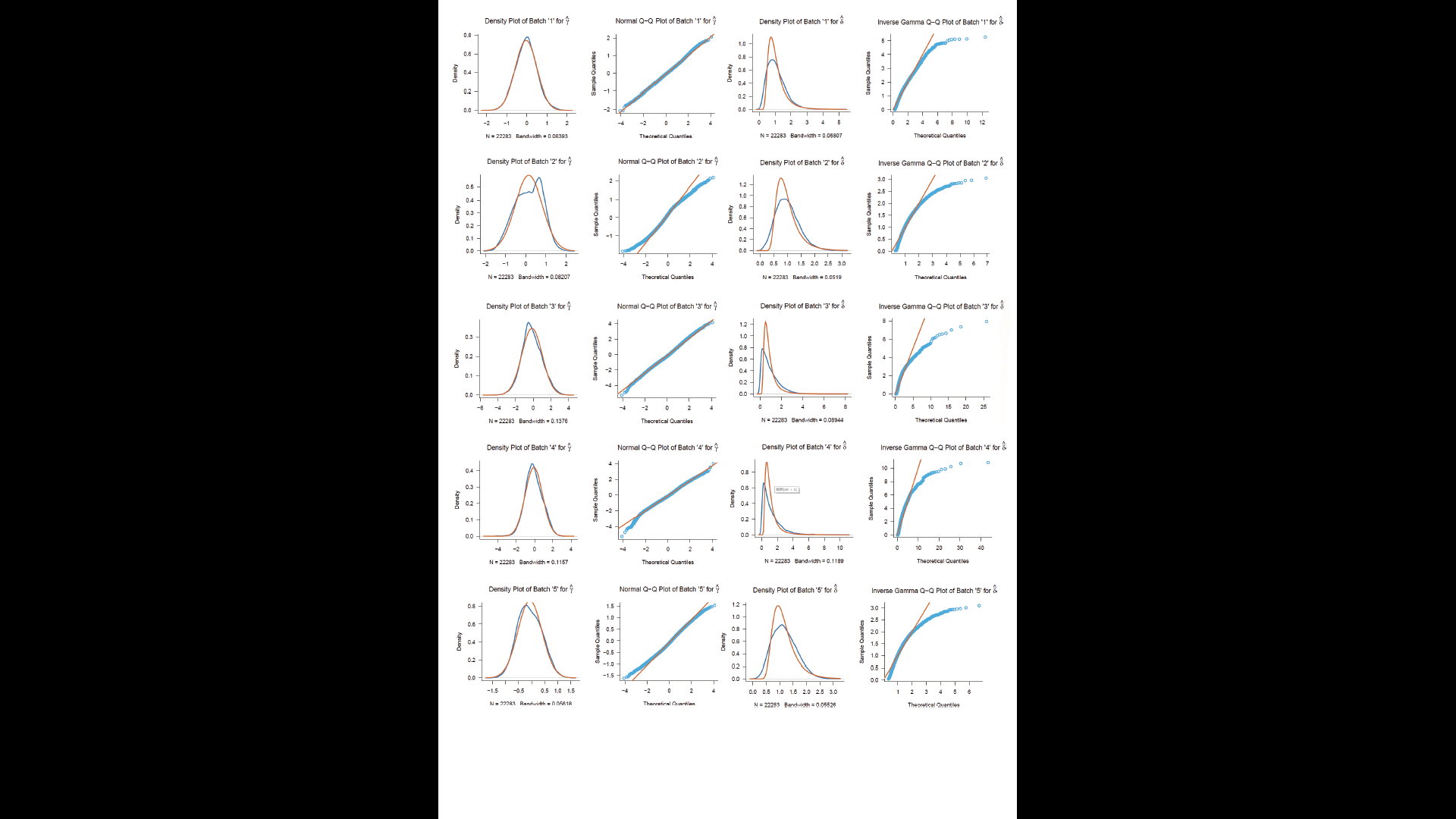

**Supplementary Figure 3. Prior plot of batch fitting effect by autoComBat by the third transcriptomic data set.**
